# Computational Analysis of Fibroblast Subpopulation Dynamics as a Driver of Fibrotic Foci Formation

**DOI:** 10.64898/2026.08.03.742640

**Authors:** Julie Leonard-Duke, David J. Csordas, Riley T. Hannan, Catherine Sano, Darius Hossainian, Mariska Batavia, Robert Andrews, Mariapaola Ambrosone, Taylor G. Eggertsen, Tania E. Velez, Jeffrey M. Sturek, Anne Sperling, Daniel Abebayehu, Thomas H. Barker, Catherine A. Bonham, Jeffrey J. Saucerman, Shayn M. Peirce

## Abstract

Fibroblasts maintain the extracellular matrix (ECM) to support tissue homeostasis and wound healing. In fibrotic diseases, fibroblasts are a primary driver of disease progression through excess collagen secretion and enhanced contractility. Replacing native tissue with a collagen rich fibrotic scar leads to a decline in tissue function. Recent research into idiopathic pulmonary fibrosis (IPF) has identified fibroblast subpopulations that may be primed for the hyper-activation that leads to increased progression of fibrotic disease. Understanding the contribution of these subpopulations to disease progression requires integrating experimental and computational techniques to understand their dynamic contributions to tissue phenotype. Herein, we introduce a framework for modeling subpopulations using a multiscale mechanistic computational model to understand differences within subpopulations, at the intracellular level and how these differences contribute to cell-and tissue-level pathology. We build and validate this framework using two well-defined subpopulations of fibroblasts in IPF. The subpopulations are defined by the presence or absence of Thy-1, a cell-surface protein that regulates fibroblast mechanosensing. We first developed a logic-based network model of a fibroblast. We then applied this model to identify sub-networks that regulate myofibroblast marker expression in the two subpopulations. Coupling this with an agent-based model (ABM) of the lung microenvironment, we observed how different rules regulating cell fate decisions in each subpopulation affected collagen content. Computational image outputs were analyzed with the open-source biological image analysis software QuPath to quantify how changes in subpopulation dynamics change model-predicted foci characteristics such as size and collagen density. We find that the ability for Thy-1^+^ fibroblasts to transition to Thy-1^-^ fibroblasts significantly increases total collagen content, as well as influences fibrotic foci characteristics. Overall, we present a combined experimental and computational framework for studying how dynamic changes in fibroblast subpopulations lead to tissue-level disease phenotypes.

**Author Summary:** In wound healing fibroblasts are responsible for rebuilding the scaffolding, called the extracellular matrix (ECM), of the damaged tissue to aid in regeneration. In fibrosis, the normal wound healing processes are hijacked leading to overproduction of ECM proteins, such as collagen, by fibroblasts leading to fibrosis. In fibrotic diseases with no known cause, such as idiopathic pulmonary fibrosis (IPF), subpopulations of fibroblasts have been identified as possible drivers of disease. Herein, we use a multiscale computational model that represents intracellular, cellular, and tissue level signaling to study how the presence or absence of a single protein on a fibroblast’s surface, Thy-1, affects the formation of fibrotic scar in IPF. Thy-1 regulates how a fibroblast senses the stiffness of the microenvironment. The absence of Thy-1 impacted intracellular signaling and when combined across many cells led to fibrosis at the tissue level. Additionally, if normal Thy-1^+^ fibroblasts could dynamically lose Thy-1 expression the amount of collagen they produced correlated with that in end stage IPF lungs. This model presents a framework for studying different fibroblast subpopulations using multiscale computational modeling to understand how changes in a single protein in one cell can, over an entire population, affect the dynamics of disease progression.

## Introduction

Cells are constantly receiving a myriad of signals from neighboring cells, the extracellular matrix (ECM), and their own transcriptional regulators. The integration of these signals within a cell determines its behavior, and the summation of a multitude of cell behaviors in a tissue leads to an emergent tissue phenotype. Disease can arise when either the external cues that cells experience become pathologic, or when internal regulatory pathways prime cells for pathological activation. Multiscale computational modeling is an ideal tool for studying this type of system. By integrating data across intracellular-, intercellular-, tissue-, and organ-level scales, models can determine how perturbations in different signals result in emergent tissue phenotypes, permitting the exploration of mechanisms that are impossible with current wet lab methods(1). Multiscale computational models have been used extensively to develop new understanding about the pathogenesis of fibrotic disease(2–6), muscle and tendon remodeling(7–9), infection(10–12), and vascular remodeling(13–15), among other processes. The present work is built upon two previous computational models developed by our group. The first of which combined intracellular logic-based signaling network modeling with an agent-based model (ABM) to explore fibroblast activation in response to different fibrotic and inflammatory cues(2). The second model was built on a multiscale framework to simulate multiple cell types at each biological scale in order to explore how matrix stiffening in idiopathic pulmonary fibrosis (IPF) affects microvascular remodeling(6). Herein, we expand upon these frameworks by simulating multiple cells in a virtual tissue cross-section to study fibrosis progression in an *in silico* model of IPF.

IPF is a progressive and fatal disease characterized by the heterogeneous formation of fibrotic lesions throughout the lung that result in a sharp decline in lung function and an average survival after diagnosis of 2-3 years(16–19). These lesions are a result of excess collagen production by myofibroblasts and increased myofibroblast contractility, which increases ECM stiffness(16, 19, 20). In measurements obtained using atomic force microscopy of mature scar tissue from the lungs of IPF patients, the stiffness of mature scar (20 kPa) has been measured to be up to ten-fold that of healthy lung tissue (2 kPa)(21). These scars form in lesions called fibrotic foci. This elevated ECM stiffness, combined with increased levels of growth factors, such as tumor growth factor beta (TGF-β), creates a pro-fibrotic microenvironment(20). The three drugs approved by the FDA to treat IPF, nintedanib, pirfenidone, and nerandomilast target several of the pathways that contribute to fibroblast-to-myofibroblast differentiation; however, they have limited impact on lung function decline and survival(22, 23). The need to create better treatments necessitates a deeper understanding of disease pathology and more advanced tools to probe the underlying biology of disease progression.

Single-cell RNA sequencing (scRNAseq) has emerged as a widely-used tool for exploring pathologic populations in IPF in recent years. Potentially pathological subpopulations of fibroblasts(24–30), macrophages(24, 28, 31, 32), and epithelial cells(24, 26, 28, 33, 34) have been identified over the years, and many can be explored using the IPF Cell Atlas(35). Subpopulations of cells are defined by differential gene or protein expression that distinguishes them from other cells of the same type. Studying these subpopulations in isolation, however, only accounts for part of the picture, and the impact of subpopulations dynamics (i.e., cells interacting with one another and with their environment) is what drives the tissue-level changes that cause the decline in organ function. Moreover, it is important to understand how different cell subpopulations and their relative abundance contribute to disease progression to develop more informed diagnostics and therapeutics. For example, what percentage of the total fibroblast population the subpopulation accounts for affects fibrosis formation.

Due to the complex fibroblast heterogeneity and multitude of mechanical and chemical cues involved in IPF, we hypothesize that the balance of different fibroblast subpopulations in the lung directly impacts the accumulation of collagen and formation of spatially heterogeneous fibrotic foci in the IPF lung. To study how fibroblast subpopulation dynamics affect fibrosis formation, we created a multiscale computational model that combines logic-based signaling network modeling to represent internal cell signaling pathways with ABM to represent cell-to-cell, cell-to-matrix, and tissue-level phenomena. We used two subpopulations of fibroblasts that were distinguished by the expression (or lack of expression) of Thy-1, a surface glycoprotein that regulates fibroblast mechanosensitivity. The absence of Thy-1 on the surface of a subset of fibroblasts has been associated with higher migration and proliferation rates and decreased apoptosis extending fibroblast lifespan (36–38). Thy-1 has also been suggested to play a role in IPF, and Thy-1 knockout mice develop markedly worse non-resolving fibrosis in response to bleomycin treatment(39). By simulating Thy-1-expressing and non-expressing fibroblasts using our multiscale model, we predict the primary responses of each subpopulation to changes in their local environments. Then, we use the model to predict how the presence, relative proportions, and origins of these subpopulations affect fibrotic lesion formation in IPF.

## Results

### Overview of multiscale model design and implementation

Our multiscale computational model was designed to represent intracellular, cellular, and population level mechanisms to predict tissue level outcomes in order to investigate the contributions of different fibroblast subpopulations to fibrotic foci formation in IPF (**Fig 1**). Based on previous studies(37, 39, 40), one differentiator of fibroblast subpopulations in the pro-inflammatory lung environment is the presence or absence of the cell surface protein Thy-1. To explore how the expression, or lack of expression, of Thy-1 affects the respective cell behaviors of Thy-1-positive (Thy-1^+^) and Thy-1-negative (Thy-1^-^) fibroblasts subpopulations, we developed a logic-based differential equation network model in Netflux(41) of intracellular signaling that includes pathways relevant to fibroblast activation and IPF that intersect with pathways regulated by Thy-1. This base model was validated against the literature and 77% of model predictions matched experimental results. To distinguish Thy-1^+^ from Thy-1^-^ fibroblasts, we set the input weight (w) for Thy-1, w_Thy-1_, at 1 or 0, respectively. In order to verify that the model would correctly predict normal lung fibroblast responses to pro-fibrotic stimuli, such as exposure to high levels of TGF-β-1 (w_TGFβ_ = 0.5) and/or environmental stiffness (w_mechanical_ = 1), for the Thy-1^+^ fibroblast (w_Thy-1_ = 1) we simulated the network model for each of these perturbations and compared it to the baseline scenario (w_TGFβ_ = 0.05, w_mechanical_ = 0.2). These model predictions were then compared with experimental findings to validate that our model correctly predicted responses to each perturbation across several key nodes in the network, including ROCK and αSMA.

**Fig 1.**
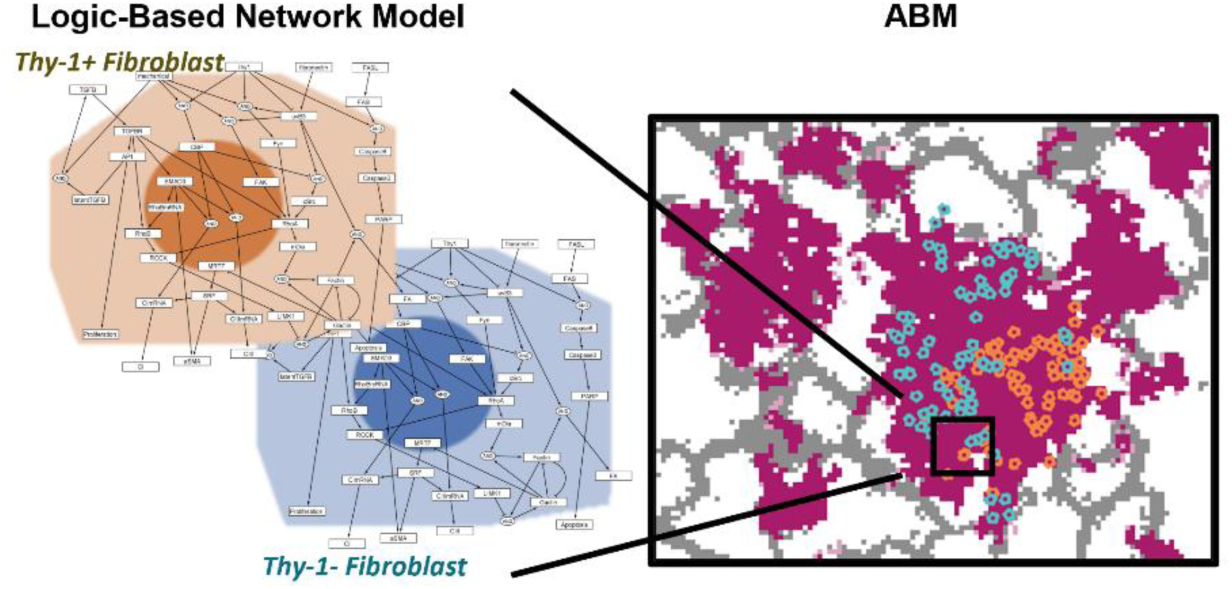
Overview of multiscale model framework. On the left is the visualization of the logic-based network model with the colors matching the color of agent in the ABM on the right. In the ABM, Thy-1^+^ fibroblasts are represented as orange agents and input w_Thy-1_ = 1 into the logic-based network model. Thy-1^-^ fibroblasts are represented by blue agents and will input w_Thy-1_ = 0 into the logic-based network model. As the agents secrete collagen the patches of the ABM will change from grey (representing the interstitial space) or white (representing the alveolar space) to pink.

Next, we connected the logic-based model to an ABM of the lung environment built in NetLogo(42) using the Py extension in NetLogo. In brief, each fibroblast agent in the ABM records the biomechanical and biochemical cues it is exposed to, such as TGF-β and ECM stiffness, normalized on a scale from 0 to 1, as described in Methods. These values, along with Thy-1 expression levels, are used as input reaction weights (w) in the logic-based network model, and the network model’s outputs are reported back to the agent, which performs the output, such as secreting collagen or undergoing apoptosis, in the ABM environment. Fibroblasts in the ABM are placed randomly in an *in silico* lung environment with a map of the interstitial versus alveolar space imported from a hematoxylin and eosin (H&E) stained histology image of a healthy human lung. As the fibroblast agents interact with the environment and with each other, they secrete collagen if stimulated by TGF-β or increased ECM stiffness. This leads to the formation of collagen-dense areas in the simulation space, representing fibrotic foci over time. Parameterization of variables that could not be identified from the literature or experimentally, such as the rates of collagen secretion and degradation, was performed using approximate Bayesian computation, as described in the Methods. We then used the multiscale model to investigate both intracellular-and population-level drivers of fibrotic foci formation and characteristics in IPF.

### Development and Validation of Lung Fibroblast Logic-Based Network Model

Logic-based network modeling has previously been used to model cardiac fibroblasts, pulmonary arterial fibroblasts, endothelial cells, and pericytes, among other cell types. In order to develop a lung specific fibroblast signaling network model that encompassed Thy-1 regulation of mechanosensing, we manually reconstructed a lung fibroblast signaling network from the literature with six signaling pathways regulated by five biochemical or biomechanical stimuli: Thy-1, TGF-β, mechanical activation, fibronectin, and FasL. Thy-1 was a direct input for four of the six pathways represented. In summary the final model contains 38 nodes that represent different mRNAs, proteins, and cell processes which are involved in 41 unique reactions(**Fig 2A**).

**Fig 2.**
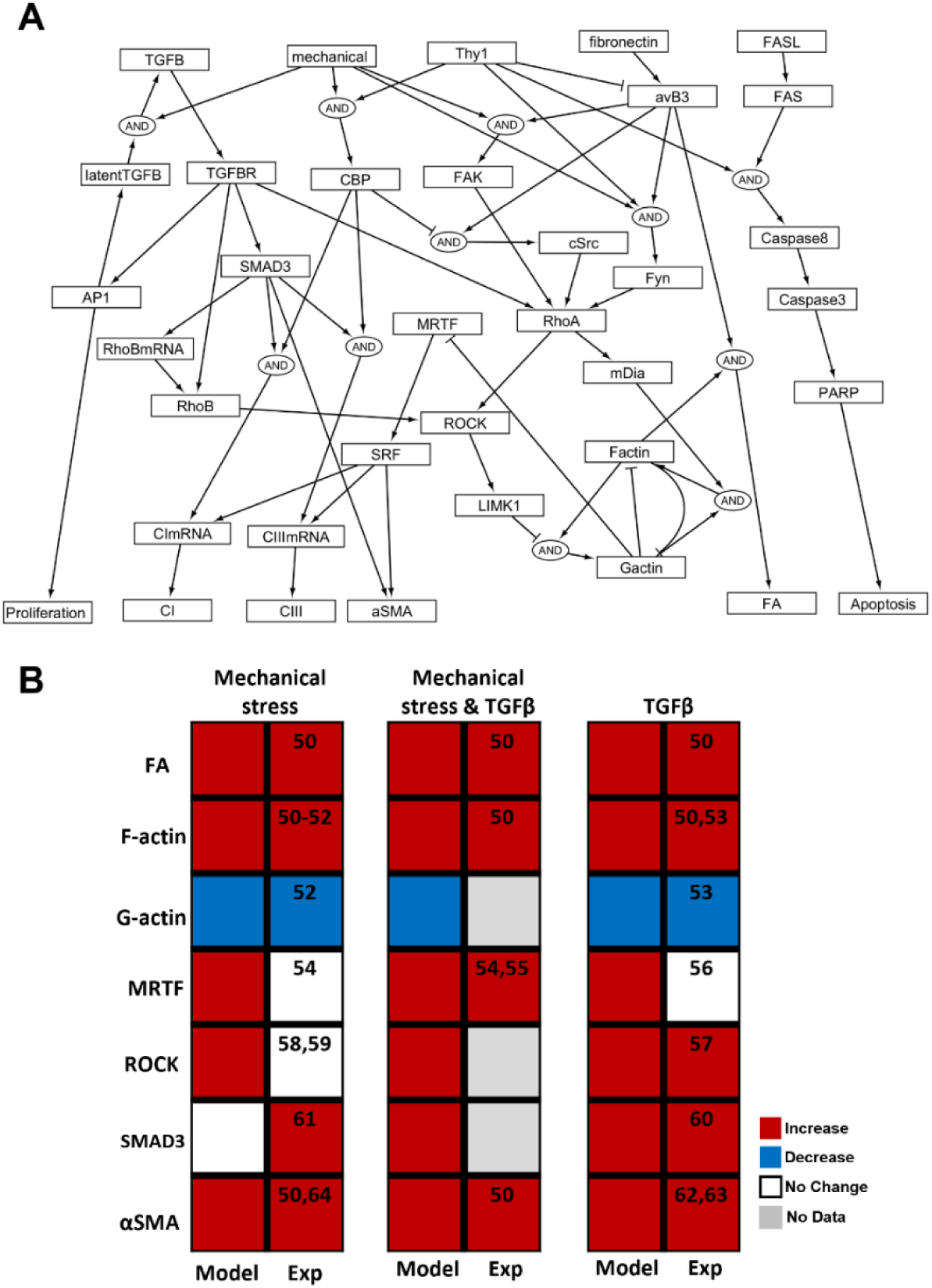
Development and Validation of Lung Fibroblast Logic-Based Network Model. (A) Logic-based network model of a lung fibroblast. (B) Validation of Thy-1^+^ normal lung fibroblast model to different perturbations relevant to the multiscale model.

To compile the validation dataset for the baseline lung fibroblast model representing normal, Thy-1^+^ fibroblasts a literature search was conducted where papers were identified using the PubMed database by searching for the perturbation (e.g., TGF-β), the output (e.g., Collagen 1), and “lung fibroblast”. Validation was performed on three primary experimental perturbations that were relevant to the overall multiscale model: mechanical stimulation, TGF-β stimulation, and combination mechanical and TGF-β stimulation. Seven nodes in the model critical for mechanoactivation were validated: Focal adhesion formation (FA)(43), F-actin(43–46), G-actin(45, 46), MRTF(47–49), ROCK(50–52), SMAD3(53, 54), and αSMA(43, 55–57). At steady state, activity in the output node after the perturbation was compared to output of the same node without perturbation. A change of more than 10% was quantified as and “increase” if the perturbation increased output activity and “decrease” if the perturbation decreased output activity. Of the nodes for which experimental data existed in the literature for, 14 of 18, or 77%, of model outputs validated the experimental result (**Fig 2B**).

### Independent experimental validation of differential responses of Thy-1^+^ and Thy-1^-^ fibroblasts to substrate stiffness

To independently validate that our model was able to predict differences in how Thy-1^+^ and Thy-1^-^ fibroblasts respond to ECM stiffening, we simulated mechanical activation in the Netflux model for both the Thy-1^+^ and Thy-1^-^ fibroblast subpopulations and compared the predictions to: 1) new experimental results that we obtained for the purpose of evaluating model outputs, such as apoptosis, F-actin, and collagen 1 production, and 2) published results related to lipid raft proteins associated with Thy-1(36). To accomplish this independent experimental validation, we simulated four cases while keeping the other nodes in the model at baseline: 1) Thy-1^+^ fibroblasts on soft matrix (w_Thy-1_ = 1, w_mechanical_ = 0.25), 2) Thy-1^+^ fibroblasts on stiff matrix (w_Thy-1_ = 1, w_mechanical_ = 1), 3) Thy-1^-^ fibroblasts on soft matrix (w_Thy-1_ = 0, w_mechanical_ = 0.25), and 4) Thy-1^-^ fibroblasts on stiff matrix (w_Thy-1_ = 0, w_mechanical_ = 1). The outputs of seven nodes for each case were compared. At steady state, activity in the output node for the Thy-1^-^ case was compared to the output of the same node in the Thy-1^+^ case for each of the four cases. Similar to the base model validation, if the node activity was increased by more than 10% compared to the Thy-1^+^ case it was quantified as “Thy-1^-^ Higher”; if the node activity was decreased by more than 10% of the Thy-1^+^ case it was quantified as a “Thy-1^-^ Lower”, and finally if the node was not more than 10% different than the Thy-1^+^ scenario then it was quantified as “No Difference”. We performed experiments to validate the model’s outputs against immunohistochemistry data. Human lung fibroblasts were cultured on either soft collagen gels or stiff tissue culture p lastic. 48 hours after seeding, they were stimulated with tumor necrosis factor alpha (TNF-α) and interleukin 1 beta (IL-1β) or vehicle control for 72 hours to match a published protocol for generating Thy-1^-^ fibroblasts (40). **Fig 3A** shows a comparison of the model output to our experimental measurements of collagen 1. The model predictions of fibroblast subpopulations behavior when cultured on a stiff matrix were validated by experimental results, which showed that both subpopulations secreted similar levels of collagen III in a soft environment, whereas in a stiff environment, Thy-1+ fibroblasts secreted more collagen III than Thy-1^-^ fibroblasts. We compared computational model results for four nodes (F-actin, collagen I, collagen III, and αSMA) with immunofluorescent results generated in our lab, and these results are summarized in **Fig 3B**. Additionally, to validate the upstream nodes of the model representing the lipid raft interactions of Thy-1 on the cell surface, we compared model outputs to data collected by Fiore et al. where they used immunoblotting to quantify lipid raft protein adhesion in Thy-1^+^ and Thy-1^-^ fibroblasts(**Fig 3C**)(36). Of all the nodes tested, 10 of 16 (62%) model outputs recapitulated the experimental results from both our data and the literature.

**Fig 3.**
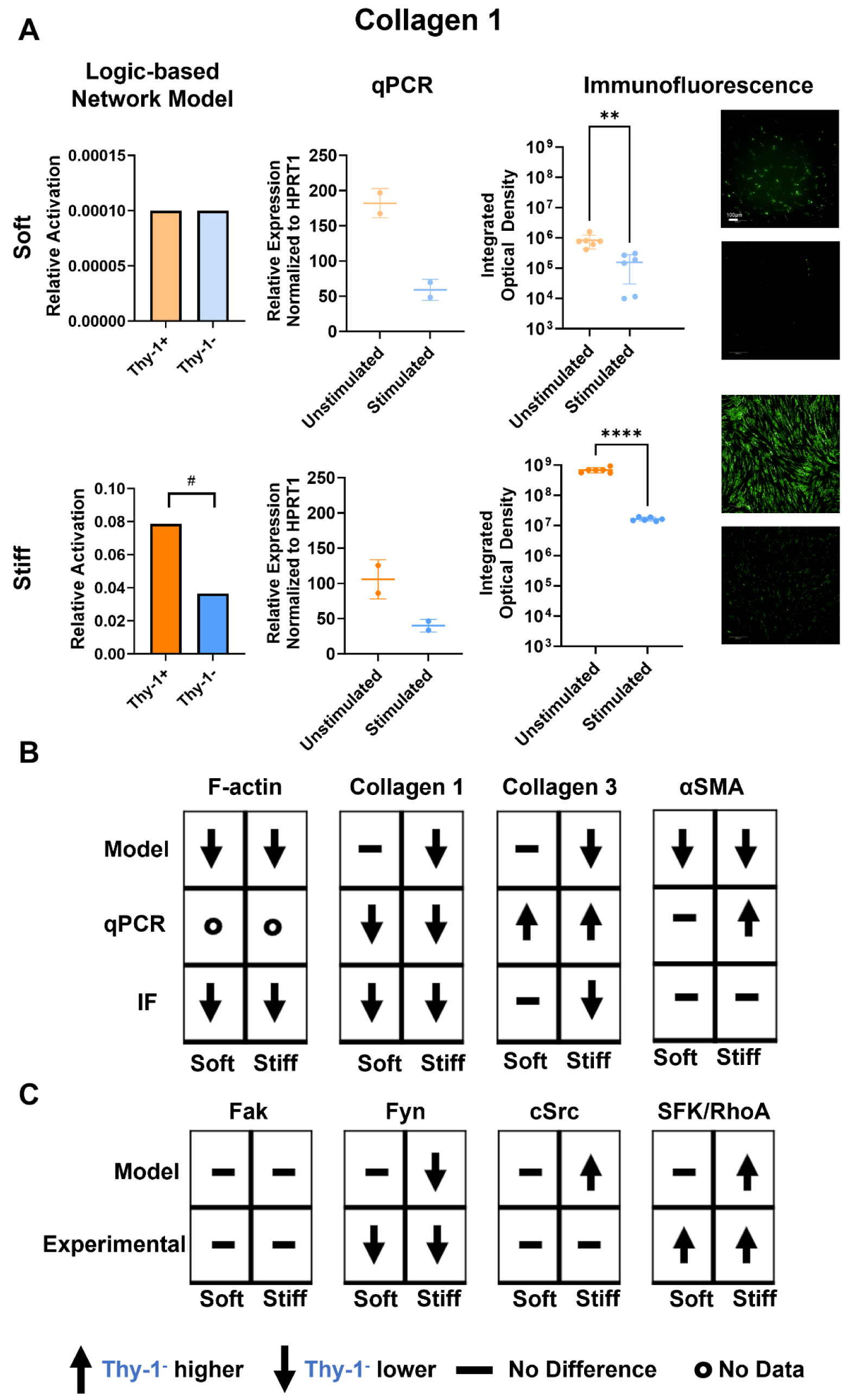
Comparison of Thy-1^+^ and Thy-1^-^ fibroblast response to hydrogel stiffness. (A) Sample validation of model output (Relative Activation) against qPCR and immunofluorescent data for collagen 1. Scale bar 100µm (B) Predicted response of Thy-1^-^ fibroblasts compared to Thy-1^+^ fibroblasts to exposure to soft and stiff environments for both computational model and experimental outputs. (C) Predicted response of Thy-1^-^ fibroblasts compared to Thy-1^+^ fibroblasts to exposure to soft and stiff environments for both computational model and experimental outputs of upstream proteins from Fiore et al(36). Statistics: For model outputs, significant differences are classified as a more than 10% change in model output (#), for immunofluorescent data, an unpaired t-test **p<0.01, ****p<0.0001

### Identifying sub-networks that regulate myofibroblast phenotype in Thy-1^-^ fibroblasts

We performed a sub-network analysis(6, 58, 59) to identify the impact of Thy-1 on myofibroblast phenotype, as defined by the expression of apoptotic markers, αSMA, and collagen 1 & 3, by comparing changes in relative activation. In the first step of the analysis, we knocked out each node in the network individually by setting its maximum activation (y_max_) to 0, then observed the impact of this knock out on apoptosis, αSMA, and collagen 1 & 3 production. We then repeated this analysis with the additional knock out of Thy-1 for each perturbation and recorded the resultant node activation for apoptosis, αSMA, and collagen 1 & 3. The nodes whose removal resulted in the largest change in the output (e.g., apoptosis) in the absence of Thy-1 were combined with the list of nodes whose function was impacted by Thy-1 knock out alone. The overlapping nodes constitute the sub-network by which the presence or absence of Thy-1 regulates myofibroblast phenotype (**Fig 4A**). This is most clearly demonstrated by the impact of Thy-1 on contributing to apoptosis, as the sub-network contains caspases 8 and 3, which directly regulate apoptosis and are expected nodes to be included in the sub-network (**Fig 4B**). Interestingly, αSMA, collagen 1, and collagen 3 node activation are all regulated by the same sub-network, which includes CBP, Fyn, and c-Src. CBP and Fyn are located on the same lipid raft as Thy-1 and therefore communicate directly with it. Additionally, CBP inhibits c-Src activity. Taken together these upstream nodes are the most important regulators of myofibroblast phenotype when comparing Thy-1^+^ and Thy-1^-^ fibroblasts.

**Fig 4.**
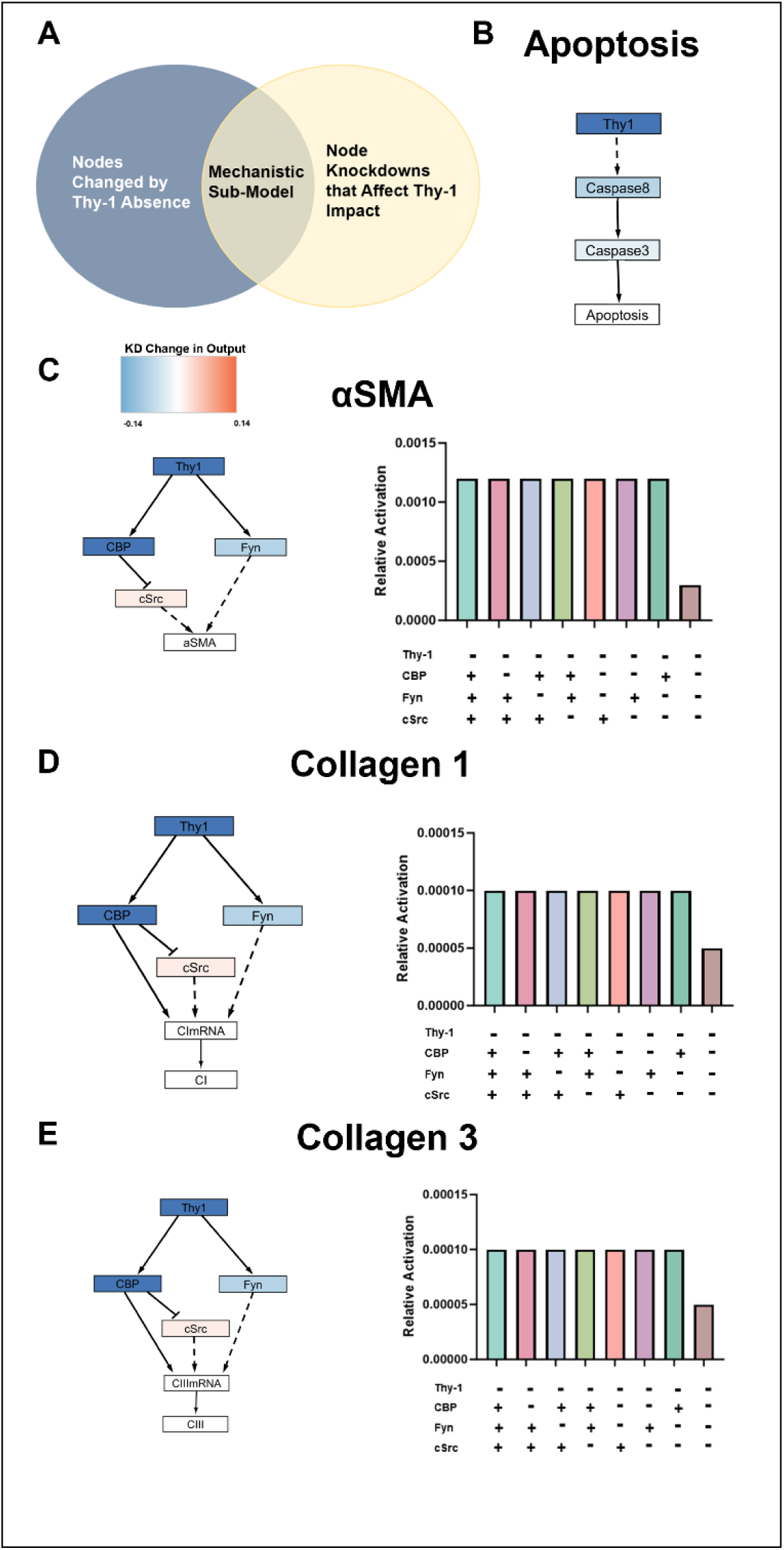
**Mechanistic sub-model analysis of Thy-1 impact on myofibroblast markers**. (A) Overview of sub-network analysis method. (B) Apoptosis sub-network. (C) αSMA sub-network. (D) Collagen 1sub-network. (E) Collagen 3 sub-network.

To validate this result, we simulated the knock out of CBP, Fyn, and c-Src individually and in combination in Thy-1^-^ fibroblasts. These lipid rafts are still activated by other inputs in the model, mechanical stimulation and αvβ3 integrin activation, but in a Thy-1^-^ fibroblast their activation is no longer regulated. As a result of this lack of regulation, all three outputs, αSMA, collagen 1, and collagen 3, were robust to single or double knock out. All three nodes had to be knocked out simultaneously in the Thy-1^-^ fibroblast scenario to see any further change in output signaling (**Fig 4C,D,&E**). This sub-network analysis highlights the importance of the lipid raft proteins in propagating mechanical signal in the absence of Thy-1 regulation.

### Effects of transient loss of Thy-1 based on fibroblast population size

To explore the dynamics that lead to the emergence of a Thy-1^-^ subpopulation, we simulated the situation where Thy-1^+^ fibroblasts lose Thy-1 after exposure to cytokines known to induce the loss of Thy-1: TNF-α and IL-1β (**Fig 5A**). For this exploration, we focused on the cell level dynamics resulting from the loss of Thy-1, not the process through which Thy-1 is lost. We defined a “transition-threshold” as a population-level variable representing the maximum allowable number of Thy-1^-^ as a percentage of total fibroblasts in the simulation. In order for a Thy-1^+^ fibroblast to transition to Thy-1^-^, the percent of Thy-1^-^ fibroblasts in the model must be below this transition-threshold. Prior studies have shown that Thy-1^-^ fibroblasts are always the smaller percentage of the total population compared with Thy-1^+^ fibroblasts, typically at most 10% of the population(39).

**Fig 5.**
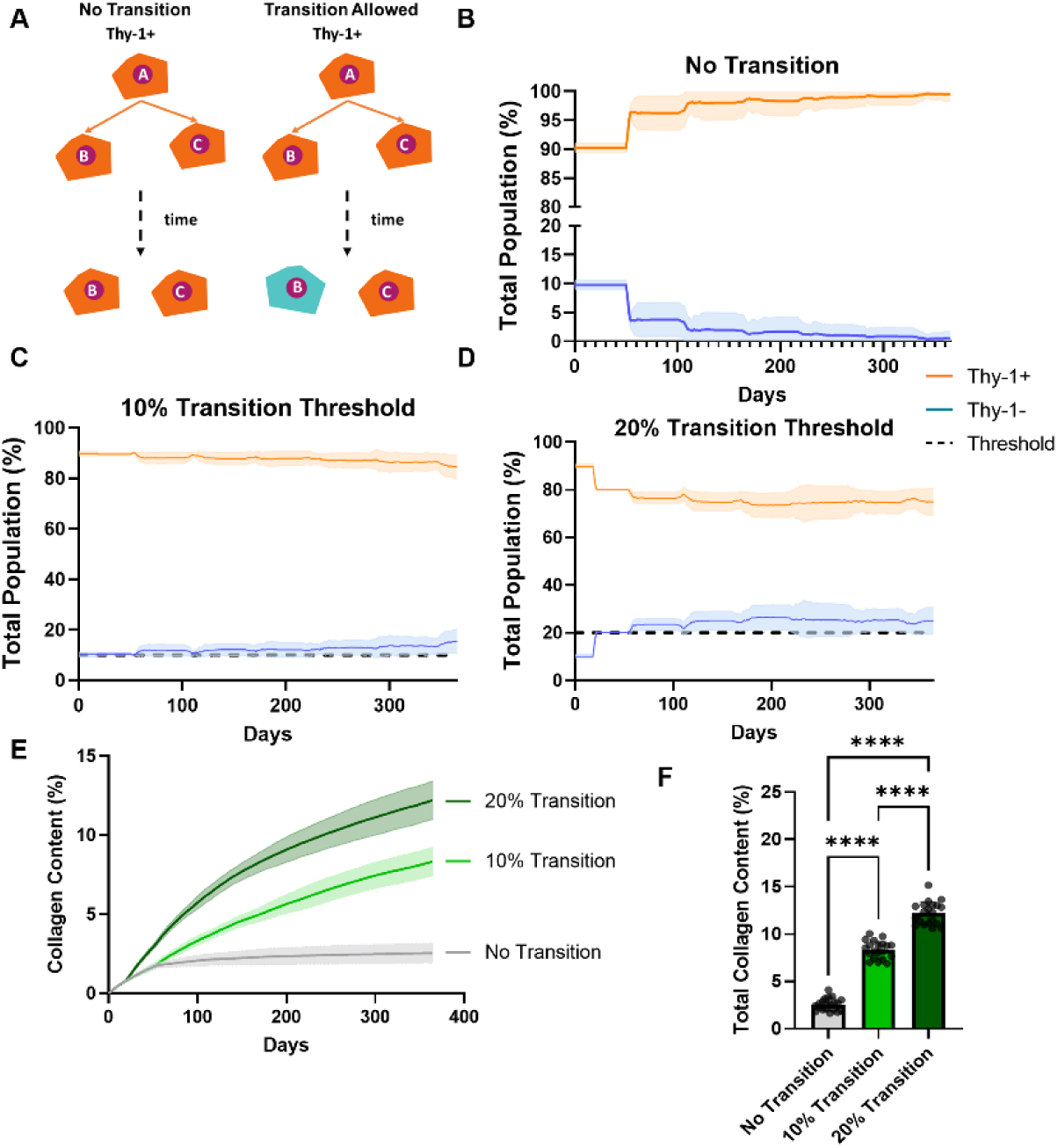
Effect of transition threshold on population dynamics and collagen accumulation in multiscale model. (A) Summary of two simulation categories. If no transition is allowed a Thy-1^+^ fibroblast (orange) and all of its progeny will remain Thy-1^+^ for the entire simulation. If transition is allowed there is a chance that the progeny of a Thy-1^+^ fibroblast will transition to Thy-1^-^(blue). (B) In the “No Transition” simulation average total population sized for Thy-1^+^ gradually increases and Thy-1^-^ decreases over the whole time course. (C) In the “10% Transition Threshold” scenario there is a steady increase in the percentage of Thy-1^-^ fibroblasts. (D) In the “20% Transition Threshold” there is a spike in Thy-1^-^ fibroblasts around Day 60 that is then maintained. (E) The total collagen content over time increases in all three scenarios, although this increase is larger in the transition permitted scenarios. (F) Quantification of total collagen content at the final time step in shows that allowing Thy-1^+^ fibroblasts to transition significantly increases total collagen content. For time course simulations: N = 20, shaded region represents standard deviation. For subfigure F, N= 20 and statistical analysis was conducted using a one-way ANOVA with Tukey’s post-hoc test. ****p<0.0001

Therefore, we set the transition-threshold to 10% or 20% to capture the beyond the full biological range of the Thy-1^-^ fibroblast. We then compared these scenarios to a scenario where Thy-1^+^ fibroblasts were prohibited from transitioning to Thy-1^-^ fibroblasts. We also assumed that Thy-1^-^ fibroblasts comprised 10% of the initial population(39).

When transitions were prohibited, Thy-1^-^ fibroblasts initiated as 10% of the population and slowly declined over time, with the average remaining slightly above zero by the end of the simulation (**Fig 5B**). When transition is permitted, Thy-1^+^ fibroblasts will transition dependent on how much IL-1β and TNF-α they have encountered in the ABM environment(40, 60). At a transition threshold of 10%, the percentage of Thy-1^-^ fibroblasts slightly increased over time (**Fig 5C**). In the 20% transition threshold scenario, there was an early initial spike in the percentage of Thy-1^-^ fibroblasts that reached steady state around Day 60. Thy-1^-^ fibroblasts comprised over 20% of the total fibroblast population for the remainder of the simulated time course (**Fig 5D**). Additionally, we evaluated how the different transition thresholds affected our primary output, collagen content, quantified as the number of pink pixels that contained notable collagen deposition to the total number of pixels. We observed an increase in collagen content over time that was different for each scenario. When Thy-1^+^ fibroblasts were prohibited from transitioning to Thy-1^-^ fibroblasts, collagen content plateaued around Day 75, and the mean collagen content remained below 5% for the entire simulation time course (**Fig 5E**). However, when Thy-1^+^ fibroblasts transitioned to Thy-1^-^ fibroblasts, collagen content increased over time. The predicted total collagen amount at the final time point, Day 365, was significantly higher when the transition threshold was set to 20% than to 10% (**Fig 5F**).

### Effect of initial subpopulations on collagen content

Prior literature suggests that few, if any, Thy-1^-^ fibroblasts reside in the healthy lung, but during disease progression the Thy-1^-^ fibroblast subpopulation emerges and expands over time(39). Therefore, we wanted to explore how the initial number of Thy-1^-^ fibroblasts affects collagen content when Thy-1^+^ fibroblasts are allowed to transition to Thy-1^-^ fibroblasts (**Fig 6A**). We used the model to predict collagen content for the three scenarios that we simulated in the prior section when the initial number of Thy-1^-^ fibroblasts was set to either 0% or 10%: 1) “No Transition”, 2) “10% Transition Threshold”, and 3) “20% Transition Threshold” (**Fig 6B**). The total collagen content at the end of one year (Day 365) was significantly higher when Thy-1^-^ fibroblasts were initially present in the lung (0% Initial: 0.23±0.05% vs. 10% Initial: 2.5±0.6%). If the fibroblasts were permitted to transition, however, the initial number did not impact the final collagen content when either the 10% transition threshold (0% Initial: 7.8±0.9% vs. 10% Initial: 8.3±0.9%) or 20% transition threshold (0% Initial: 12.2±1.1% vs. 10% Initial: 12.2±1.2%) were imposed.

**Fig 6.**
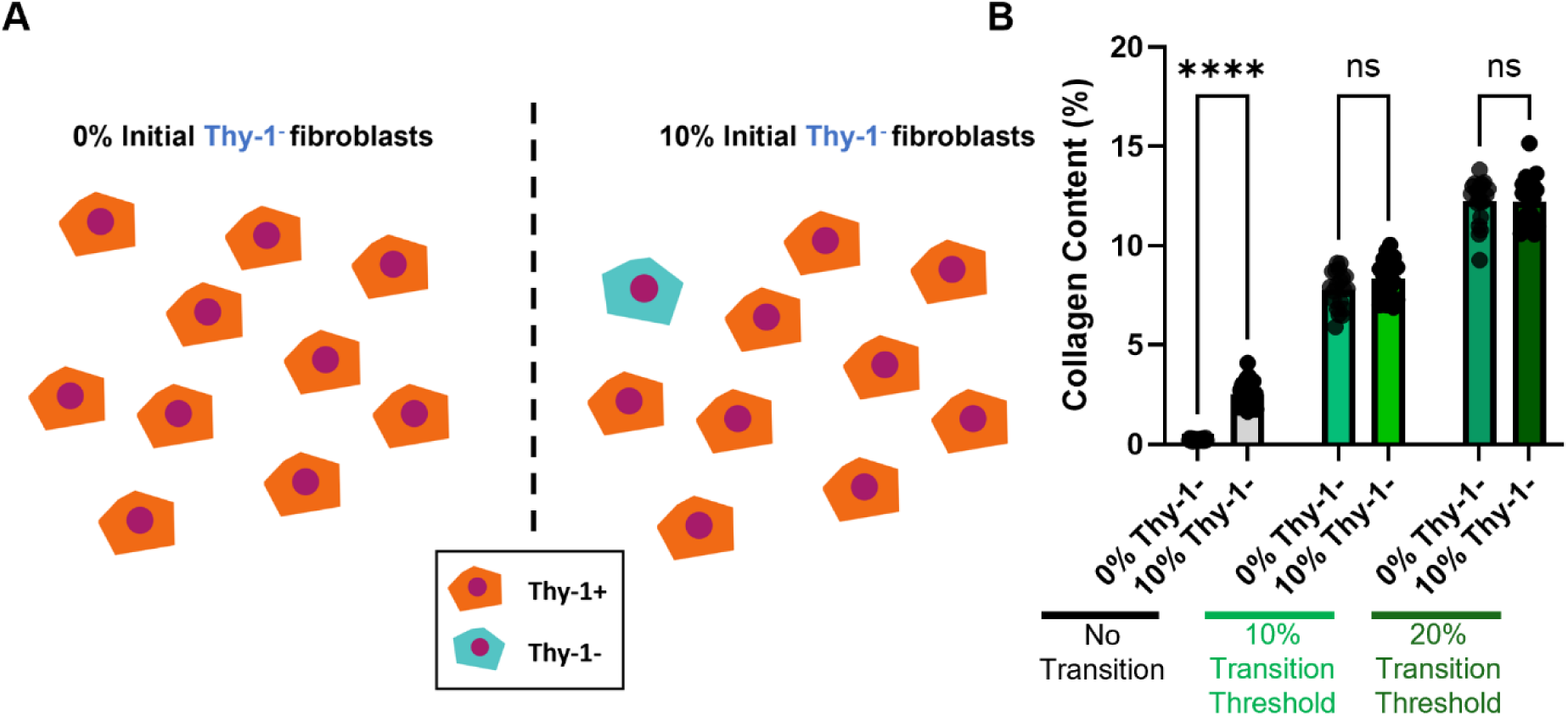
Effect of initial Thy-1^-^ fibroblast population level on collagen content. (A) Two initial subpopulation scenarios modeled based on prior literature. (B) The initial number of Thy-1^-^ fibroblasts had a significant effect on collagen content in the scenario where transitioning was prohibited; however, it did not significantly impact collagen content in the 10% or 20% transition threshold scenarios. N = 20 simulations, statistical analysis was performed with a two-way ANOVA with Tukey’s post-hoc test. ****p<0.0001, ns = not significant

### Quantification of collagen content in human IPF lung samples

To validate that our predicted collagen content levels were within a reasonable range for IPF we analyzed pentachrome stained human lung sections from IPF and non-IPF patients. A QuPath pixel classifier algorithm was trained to identify pixels positive for the collagen stain in (**Fig 7A**). The algorithm was trained and executed in a blinded fashion and twenty regions of interest (ROI), that were the same size as the computational area modeled in the ABM, were randomly selected for each sample. These twenty ROIs were then averaged per sample to account for variability. Four IPF and four non-IPF tissue samples were used in this analysis. The IPF lung sections had significantly higher collagen content, 15.3±4.9%, than the non-IPF lung sections, 3.5±3.8% (**Fig 7B**). Therefore, our computational model predictions that allow Thy^-^ transition to occur fall within the predicted range of collagen content values (10.4% to 20.2%).

**Fig 7.**
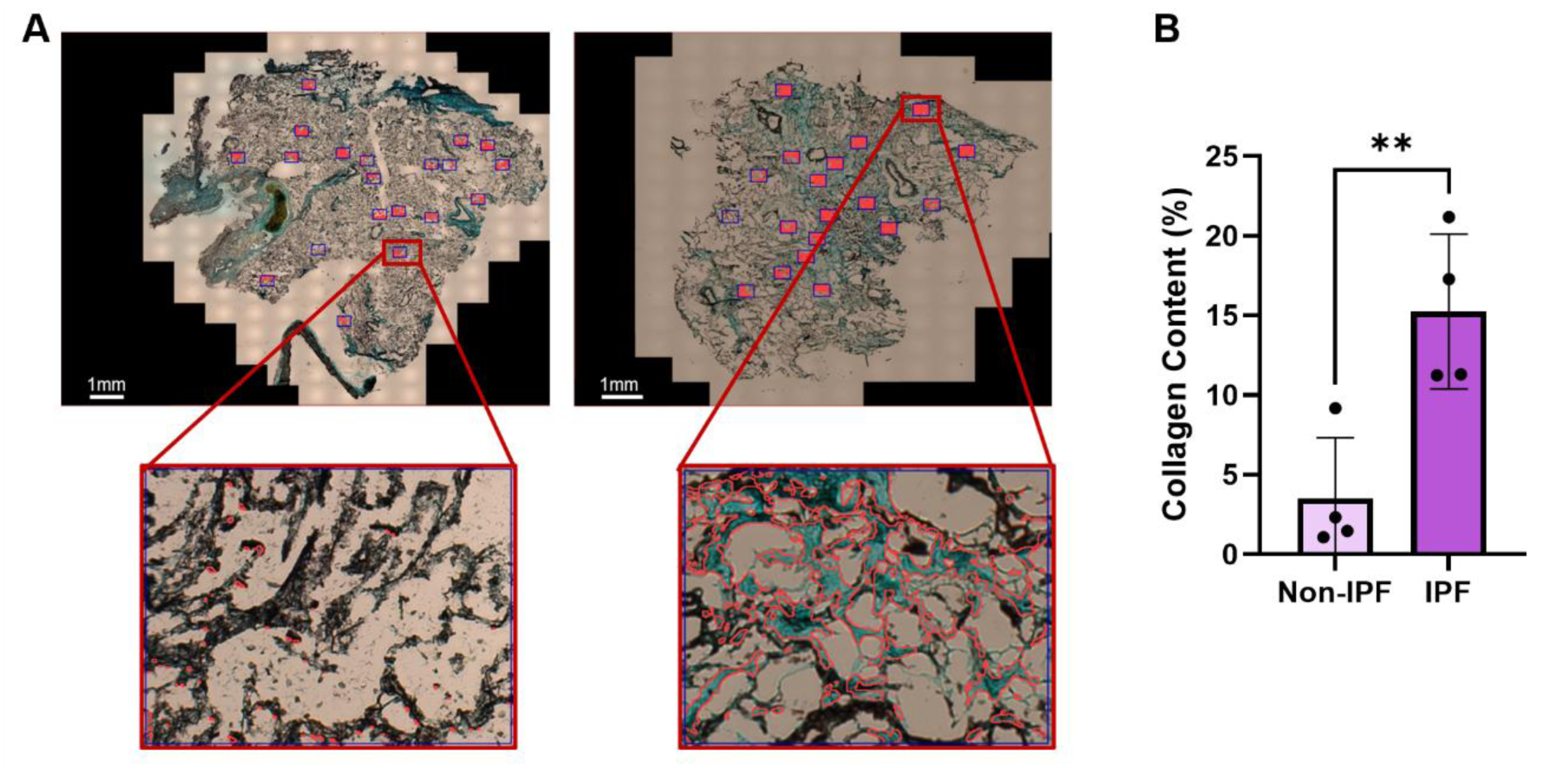
Analysis of collagen content in human lung samples with and without IPF. (A) Sample view of QuPath analysis with collagen positive pixels (blue) outlined in red. Below a zoom in on a single ROI for a non-IPF and IPF sample. (B) Total collagen content was significantly higher in the IPF samples when compared to the non-IPF controls. Statistics: unpaired two-tailed t-test, **=p<0.01.

When comparing final collagen content predicted by the model to actual pentachrome stained patient lung tissue sections only the 20% transition threshold scenario (**Fig 5F**, 12.2±1.2%) fell within one standard deviation of the average collagen content in IPF patient tissue (**Fig 7B**, 15.3±4.9%). This relationship also was true for the scenario where there are no Thy-1^-^ fibroblasts in the initialization of the model (**Fig 6B**, 12.2±1.1%). Notably, the no transition allowed scenario (**Fig 5B**, 0.2±0.05%) also fell within one standard deviation of the non-IPF collagen content, (**Fig 7B**, 3.5±3.8%) even when there were Thy-1^-^ fibroblasts initially (**Fig 5F**, 2.5±0.6%). Together this suggests that the presence of a subpopulation in healthy tissue is not as impactful as the ability for fibroblasts to transition into different subpopulations and the accumulation of these subpopulations over time.

### Fibroblast subpopulation effects on fibrotic foci patterning

In order to explore if subpopulation dynamics had an effect on the characteristics of the fibrotic foci themselves, we developed an automated image analysis pipeline in the open-source software QuPath to identify fibrotic foci in our graphical simulation outputs based on grayscale color values that represented collagen content. Our pipeline quantified characteristic spatial features of foci, such as average number of foci, average brightness per foci, average size characteristics (e.g. area and perimeter), solidity, and circularity (**Fig 8A**). Twenty simulation outputs per scenario, per time point, were used for the analysis, and all foci in the graphical simulation outputs were averaged for downstream analysis. In this manner, we identified characteristics that distinguished each graphical simulation output based on the transition condition scenario by which it was generated.

**Fig 8.**
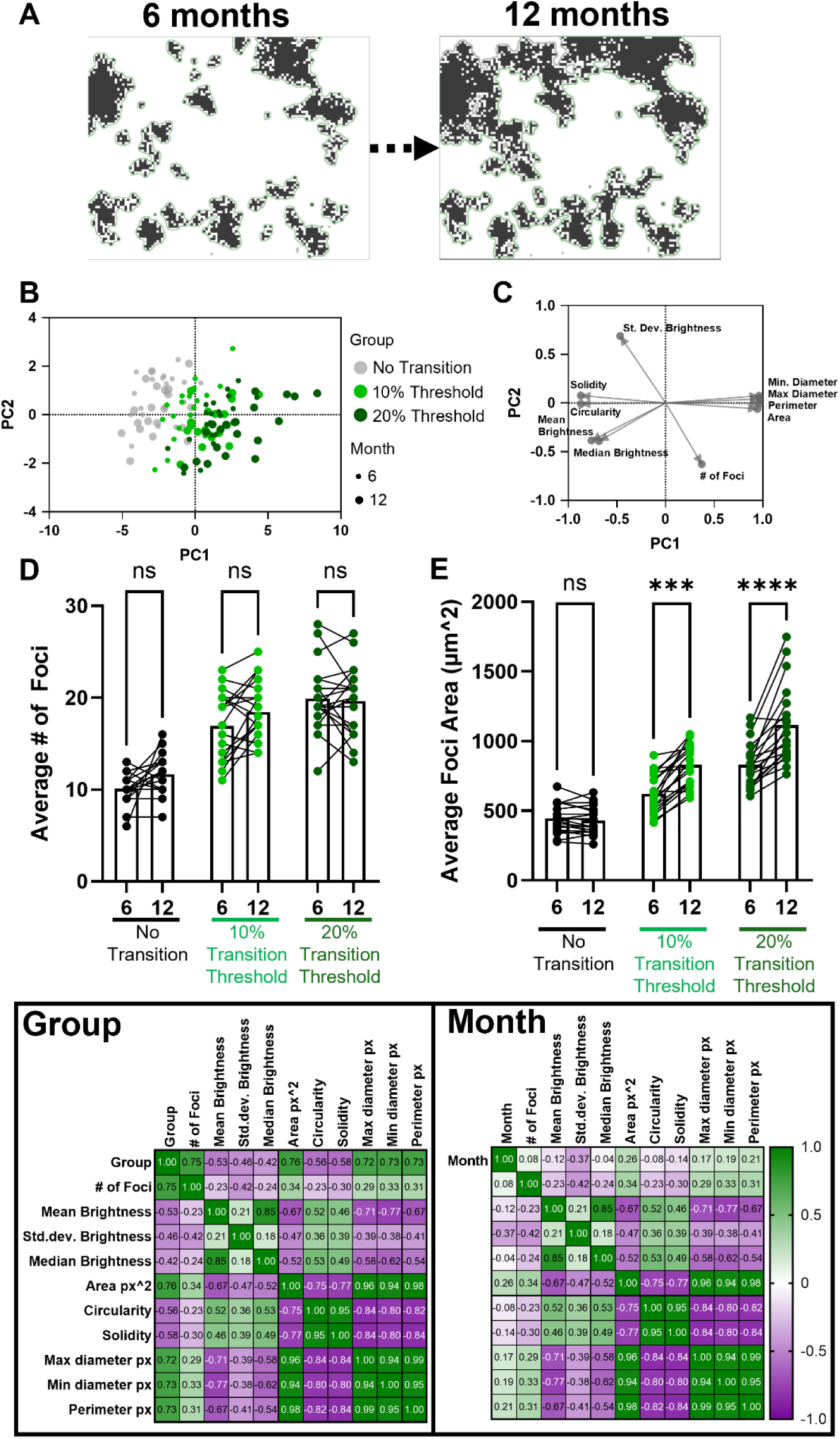
Characterization of fibrotic foci in graphical simulation outputs based on time and transition scenario. (A) Representative time course images of foci at 6 and 12 months in the 20% transition threshold scenario, with green outline representing QuPath’s identification of foci. (B) Principal component analysis of foci characteristics shows that the ability to transition strongly influences principle component 1. (C) Loadings of principal component analysis demonstrate separation between characteristics related to foci size (minimum & maximum diameter, area, and perimeter) from other characteristics such as number of foci or strength of collagen positive signal (brightness). (D) The number of foci over time did not significantly change in any of the simulation scenarios. (E) Overall foci area increased with time, with the most dramatic effect being observed in the 20% transition threshold scenario. (F) Pearson correlation matrices for the correlation between transition threshold scenario (No transition, 10% transition threshold, 20% transition threshold) and time point (6 or 12 months). Statistics: D&E: Two-way ANOVA with Tukey’s post hoc, ns = not significant, ***p<0.001, ****p<0.0001.

Principal component analysis demonstrated a separation along principal component 1 (66% of total variance) between allowing fibroblasts to transition from the Thy-1^+^ to the Thy-1^-^ subpopulation versus no transition (**Fig 8B**). Although some outputs from the 10% transition threshold scenario at 12 months clustered with those from the 20% transition threshold scenario. Analyzing the underlying loadings, we observed that metrics related to foci size, such as area, perimeter, and diameter, were positively correlated with principal component 1, whereas metrics related to shape, such as solidity and circularity, were negatively correlated (**Fig 8C**). The standard deviation of brightness within a focus and the number of foci were the inputs most related to principal component 2, which captured an additional 12% of the variance. Thus, the ability to transition contributes to larger, more irregularly shaped foci, but is not strongly correlated with the number of foci. These relationships were revealed by plotting the average number of foci, which remained constant from 6 to 12 months across all scenarios (**Fig 8D**). In contrast, the average area of each focus increased in the scenarios where Thy-1^+^ fibroblasts were allowed to transition to Thy-1^-^ fibroblasts. For the 10% transition threshold, foci area at 6 months was 622.3±137.2 square microns and increased to 828.7±148.5 square microns by 12 months. For the 20% transition threshold, foci area at 6 months was predicted to be 829.3±156.2 square microns versus 1114±274.4 square microns at 12 months. The average foci area did not change over these time points when fibroblast transitioning was prohibited (**Fig 8E**). Finally, we wanted to quantify the strength of the correlation between transition threshold and fibrotic foci characteristics. We observed a positive correlation between the level of the transition threshold (10% or 20%) and the number of foci (0.75), foci area (0.76), perimeter (0.73), and maximum (0.73) and minimum (0.73) diameter. As a control, we grouped the computational output by month (6 vs. 12), instead of the transition group, and found that timepoint alone did not significantly correlate with foci characteristics, indicating that the characteristics are more attributed to the underlying fibroblast subpopulation dynamics and not time course (**Fig 8F**).

## Discussion

As transcriptomic platforms that allow researchers to identify subpopulations of cells become more accessible so does the need for tools that allow us to interrogate the functional role of these subpopulations in disease progression. Here we presented a framework for building a multiscale computational model from subpopulation data to study the intracellular signaling pathways that differ between subpopulations and how the population dynamics of the subpopulations affect disease progression. We applied this framework to two subpopulations that are well studied in the context of IPF, Thy-1^+^ and Thy-1^-^ fibroblasts and compared computational results to *in vitro* and patient data at multiple length scales. We first studied the effect of the loss of Thy-1 on myofibroblast marker expression in a single cell using logic-based network modeling and mechanistic sub-network modeling. We then used a multiscale computational model that integrates the logic-based network model with an ABM to study how varying percentages of each population drive fibrotic foci formation over time.

Thy-1 exists on the cell surface on a lipid raft in a cluster with other proteins including CBP, Fyn, and c-Src(37). When we evaluate the mechanistic sub-network that drives the production of myofibroblast markers αSMA, collagen 1, and collagen 3, we observe that all three sub-networks are regulated by changes in activation to CBP, Fyn, and c-Src. The activation of these nodes is the main differentiator between Thy-1^+^ and Thy-1^-^ fibroblasts when regulating myofibroblast marker expression. Importantly, Thy-1^-^ fibroblasts were robust against additional knock outs of one or two of these markers in combination with the loss of Thy-1 indicating some parallelism in the production pathway. Only when all three nodes were knocked out in combination with Thy-1 loss was myofibroblast marker activation further decreased. This highlights the importance of Thy-1 regulation of its co-lipid raft proteins in sensing mechanical stimuli.

One advantage of computational models is that they enable the exploration of different cell subpopulation dynamics that cannot easily be studied using *in vitro* or *in vivo* experimentation. We applied our multiscale computational model to study and compare how: 1) the initial proportion of fibroblast subpopulations (0% Thy-1^-^/100% Thy-1^+^ vs. 10% Thy-1^-^/90% Thy-1^+^) in the lung, and 2) the dynamic transition of Thy-1^+^ fibroblasts to Thy-1^-^ fibroblasts affected collagen content over time. The ability for Thy-1^+^ fibroblasts to transition and the population level rules that dictated this transition significantly impacted population dynamics over time and final collagen content.

The scenario allowing Thy-1^+^ fibroblasts to transition with a transition threshold of 20% was the most closely approximated by the independent experimental data and captured more biological variability, as reflected by larger standard deviations for both the percentages of each subpopulation over time and total collagen content over time. IPF is a highly heterogenous disease, both between patients and within different lung regions of the same patient. Therefore, having a model that can capture a wide range of pathological outcomes will be helpful in the future for identifying putative pharmacological perturbations that are effective in and across different stages of disease progression.

To take a step beyond analyzing the overall collagen content predicted by our model for the different scenarios, we developed an analysis pipeline in QuPath that could identify foci and then characterize their size, brightness, circularity, and solidity. A principal component analysis demonstrated that the ability to transition (10% and 20% transition thresholds) led to similar fibrotic foci characteristics when compared to no transition. This effect was more apparent at the twelve-month time point. The number of foci did not significantly change in any of the simulation scenarios, but the average size of the foci significantly increased in both the 10% and 20% transition threshold scenarios, indicating that the expansion of existing fibrotic niches was the main driver of collagen accumulation as opposed to the formation of new foci. A correlation analysis confirmed that the most influential differentiator between simulation scenarios was the number of foci and foci geometry characteristics.

Our multiscale modeling approach has the potential to enable future studies that explore the impact of fibroblast subpopulation dynamics on fibrotic foci formation in IPF using publicly available data sets. We used a well-defined subpopulation in IPF that had an easily identifiable surface marker, Thy-1, to demonstrate how experimental and computational techniques can be integrated to dynamically subpopulations(36–40, 60, 61). We would hope to extend this framework to include other fibroblast subpopulations that are evident from scRNAseq datasets. A recent study by Tsukui et al. identified one population that was exclusively present in fibrotic lungs was characterized by the expression of collagen triple helix repeat containing 1 (Cthrc1) through scRNAseq analysis of both bleomycin-treated Col1a1-EGFP mouse and lung samples from patients with IPF, scleroderma, and healthy controls(29). This Cthrc1^+^ subpopulation was found to express the highest levels of collagen and was significantly more migratory than other lung fibroblast subpopulations, suggesting it could be a driver of fibrosis progression even though it represents only a subset of fibroblasts in a fibrotic lung(29). In future extensions of this model, the impact of other signals in the microenvironment, such as proinflammatory signals, should be incorporated to simulate how different subpopulations respond to paracrine cues from other cell types, such as macrophages and epithelial cells(62, 63). Lastly, this model could be extended to incorporate other related processes that affect the progression of IPF and response to therapeutics such as angiogenesis and microvascular regression(6).

In conclusion, multiscale models are a powerful tool for data integration across biological scales that have the potential to advance our understanding by predicting dynamic tissue changes that cannot be studied *in vivo*. To our knowledge, our multiscale model is the first to study fibroblast subpopulations in IPF using an ABM to represent the tissue level consequences of cell subpopulation dynamics. Ultimately, these models can be used to identify potential therapeutic approaches that target cell subpopulations to enhance the efficacy of existing IPF treatments.

## Methods

### Logic-Based Model

A lung fibroblast signaling network encompassing pathways regulating fibroblast-to-myofibroblast activation, as well as Thy-1-mediated mechanosensing, was manually reconstructed from the literature. In summary, six signaling pathways regulated by five biochemical or biomechanical stimuli: Thy-1, TGF-β, mechanical activation, fibronectin, and FasL. Thy-1 was a direct input for four of the six pathways represented.

A literature review was conducted to identify the role of the pathways above in fibroblasts with a focus on Thy-1^-^ fibroblasts. A subset of the papers that reported studies in which TGF-β stimulation, mechanical stimulation, or a combination of the two was performed was set aside for validation of the Thy-1^+^ normal fibroblast model. A second subset of studies that directly described Thy-1 regulation of lung fibroblast response to the identified stimuli above were used for model development. All reactions identified as being regulated by Thy-1 were built from studies that used lung fibroblasts from mouse, rat, or human sources. The final model contains 38 nodes that represent different mRNAs, proteins, and cell processes which are involved in 41 unique reactions.

In order to convert the fibroblast signaling network diagram into a logic-based ordinary differential equation (ODE) based computational model, we utilized the software “Netflux”, as previously described(41, 64). In summary, a normalized Hill ODE is implemented to represent the activity of each node according to default parameters and logic gating. For each species (mRNA, protein, cell process) the default parameters are set to include an initial concentration of 0 (y_int_), maximum activation of the species at 1 (y_max_), and time constant ꞇ. The exception to the default value for initial is G-actin which we assume there to be the maximum amount available (y_int_ = 1) prior to stimulation which is subsequently consumed to form F-actin. The time constant parameter ꞇ is scaled for each species according to the type of reaction it is involved in with signaling reactions requiring 6 minutes, transcription reactions requiring 1 hour, and translation reactions requiring 10 hours. The reactions in the model also contain three parameters: reaction weight (w = 1), the Hill coefficient (n = 1.4), and EC50 (EC50 = 0.6). The Excel sheet containing the constructed lung fibroblast model can be found in the Supplementary Materials. The final fibroblast network was visualized using Cytoscape(65).

### Validation and Sensitivity Analysis of Logic-Based Network Model

To validate the baseline fibroblast model representing normal, Thy-1^+^ fibroblasts a literature search was conducted where papers were identified by searching for the desired perturbation (e.g., TGF-β), the desired output (e.g., Collagen 1), and “lung fibroblast” into the PubMed database search query. Three experimental perturbations that were relevant to the overall multiscale model were used for validation: mechanical stimulation, TGF-β stimulation, and combination mechanical and TGF-β stimulation. Seven nodes in the model critical for mechanoactivation were validated: Focal adhesion formation (FA)(43), F-actin(43–46), G-actin(45, 46), MRTF(47–49), ROCK(50–52), SMAD3(53, 54), and αSMA(43, 55–57). At steady state, activity in the output node after the perturbation was compared to output of the same node without perturbation and if the node activity was more than 10% higher than baseline it was quantified as an “Increase”, if the node activity was more than 10% lower than baseline it was quantified as a “Decrease”, and finally if the node was not more than 10% different from baseline then it was quantified as “No Change”. Of the nodes for which experimental data existed in the literature for, 14 of 18, or 77%, of model outputs validated the experimental result.

### Evaluating Sub-Networks that Regulate Thy-1 Control of Myofibroblast Phenotype

To identify the nodes that regulate the effect of Thy-1 on myofibroblast markers, αSMA, collagen 1, and collagen 3, we performed a sensitivity analysis in the presence and absence of Thy-1. Knocking down each node in the absence of Thy-1 describes the role of that node in Thy-1^-^ fibroblasts. Comparing the difference in node activity in the presence of Thy-1 compared to the activity in the absence of Thy-1 describes the role of that node in regulating Thy-1’s effect on myofibroblast phenotype. The nodes that have a differential effect dependent on Thy-1 are identified as the sub-network that can be used as a mechanistic description of Thy-1 activity. The sub-network analysis is performed in MATLAB then visualized used Cytoscape(65).

### Agent-Based Model (ABM)

The ABM portion of the model was built in NetLogo(42), a freely available ABM software, and is a 2D representation of a 400μm x 300μm x10 μm-thick slice of lung tissue. The 2D ABM simulation space is discretized into a square (x,y) grid of square pixels, each representing a 10μm x 10μm. An image of healthy lung histology was imported into NetLogo with the interstitial space represented by grey pixels and the alveolar space represented by white pixels. Growth factors, such as TGF-β, and cytokines, such as IL-1β and TNF-α, were equally distributed across all patches in the model and their respective concentrations were represented in pg/mL. Additionally, all patches started at the average healthy lung stiffness of 2 kPa(21).

Upon initialization, fifty fibroblasts were randomly placed within the designated interstitial space. The fibroblast-specific logic-based network model was used to dictate the behaviors of simulated fibroblast “agents”. At each time step, each agent recorded its environmental cues (i.e., stiffness, TGF-β concentration, etc.) and used them as the inputs to a logic-based network model, which in turn output cell behaviors (i.e., secrete collagen, migrate, apoptose, etc.) based on the information. The agent then executed that decision, and the process repeated on the next timestep. Each time step in the model represented 24 hours, and the total simulated time was 1 year, which is similar to previous publications from our lab and computationally tractable(20, 66, 67).

Based on previous data in literature, the initial Thy-1^-^ population ranges from 0 to 10% of the total initial fibroblast population(39). Extended exposure to TNF-α and IL-1β has been demonstrated as a potential trigger for the loss of Thy-1 expression in human lung fibroblasts(39, 40, 60). In these studies, up to 40% of fibroblasts lost Thy-1 expression when cultured with TNF-α and IL-1β for one week(40, 60). The increase in Thy-1^-^ fibroblasts could be due to just the expansion of the Thy-1^-^ population itself or also due to the loss of Thy-1 in Thy-1^+^ fibroblasts. To explore the effects of both of these scenarios we either allowed Thy-1^+^ fibroblasts to transition to Thy-1^-^ after extended exposure to TNF-α and IL-1β or prevented transition such that once an agent is assigned as Thy-1^+^ or Thy-1^-^ it and all of its progeny maintain that phenotype. It was also found that the fibroblasts that did not lose Thy-1 expression during the seven-day culture with TNF-α and IL-1β never lost Thy-1 expression, even after repeated exposure(40, 60). Because the mechanism for this resistance to Thy-1 loss is not fully understood, to incorporate the effects of it in the model it was represented as a “transition threshold”, if more than a certain percentage of the fibroblast population is Thy-1^-^ then Thy-1^+^ fibroblasts would not transition to Thy-1^-^.

### Connecting the Models Across Scales

The logic-based network model and the ABM were connected using principles from previously published multiscale models from our lab(2, 6) (**Fig 9**). The ABM contains the extracellular components of the model, including ECM stiffness, fibronectin, TNF-α, IL-1β, and TGF-β. Each patch in the ABM contains these components, and the fibroblast agent interacts with them as it migrates over them. Each time step in the coupled model represents 24 hours, and the model was run for 365 ticks (1 year). At each time step, each agent recorded the values of the biomechanical and biochemical cues present on the patch it was on, input these values into the network model, and based on the network model’s output, proliferated, underwent apoptosis, or secreted collagen. Fibroblast agents migrated randomly at every time step, and Thy-1^-^ fibroblast agents had a random chance of entering the alveolar space if the combined stiffness of the surrounding patches was above a threshold *T_a_*. Agent migration was constrained to the borders of the simulated space. Values from the ABM, which are stored in concentrations or approximate stiffness (kPa), were translated into normalized values for the reaction weight, *w*, (from 0 to 1) for the logic-based network model and represented 0 to 100% of that node activation. All initial input weights except Thy-1 and TGF-β are kept at normalized values listed in **Table 1**. The input for Thy-1 depends on if the fibroblast is Thy-1^+^ (Thy-1 = 1) or Thy-1^-^ (Thy-1 = 0). The relative activation of TGF-β receptor is described using a Hill equation where the dissociation constant K_d_ represents the ligand concentration at which 50% of the TGF-β receptors are activated at equilibrium, as we have previously described(2).

**Fig 9.**
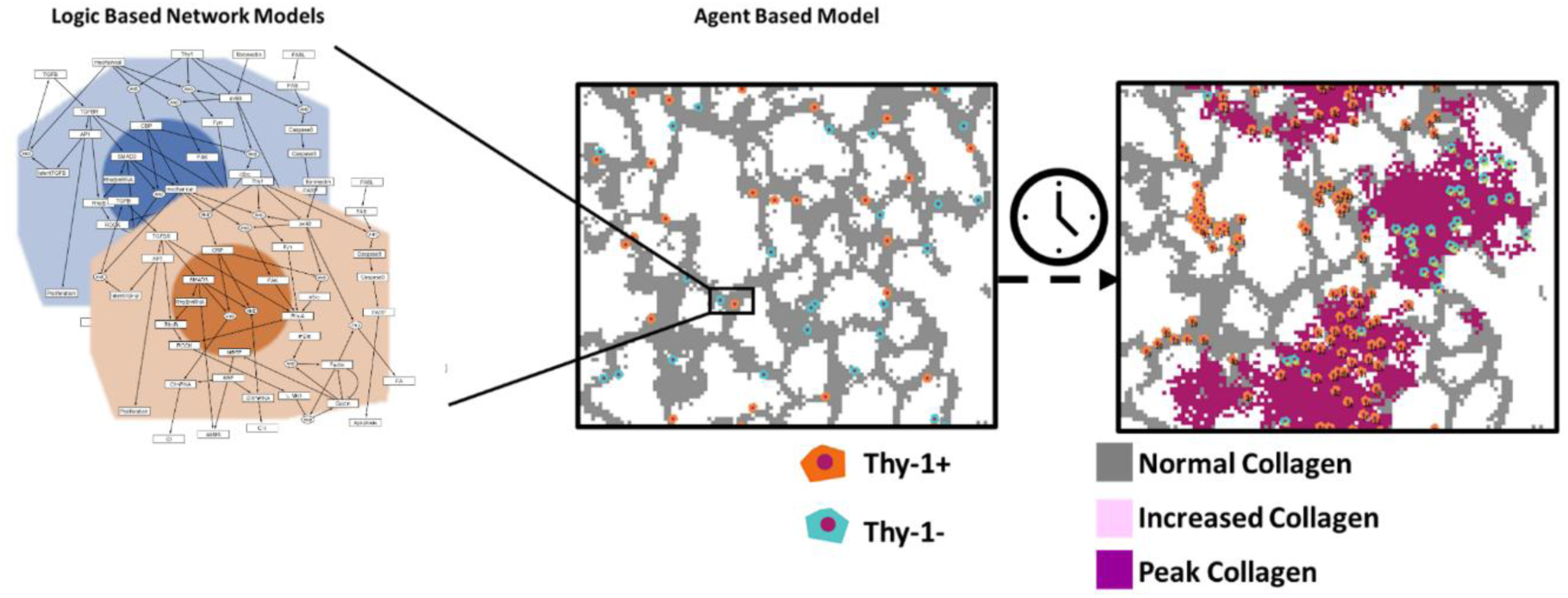
Overview of multiscale model structure. Representation of model input across scales and progression of fibrotic foci formation.

**Table 1.** Input parameters for multiscale model.

| Variable | Input Value |
| --- | --- |
| $w_{fibronectin}$ | 1 |
| $w_{FASL}$ | 0.25 |
| $w_{mechanical}(initial)$ | 0.1 |
| $TGF - \beta_{ABM}$ | 700 pg/mL |
| $K_{D,TGF-\beta}$ | 28 pM |
| $IL - 1\beta_{ABM}$ | 8750 pg/mL |
| $K_{D,IL-1\beta}$ | 500 pM |
| $TNF - \alpha_{ABM}$ | 323 pg/mL |
| $K_{D,TNF-\alpha}$ | 19 pM |

This equation is listed below in **Equation 1**, and values for TGF-β_ABM_ and K_D,TGF-β_ are listed in **Table 1:**

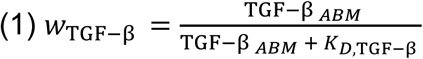

As fibroblasts secreted collagen into ECM the stiffness of the ECM increased from an initial stiffness of 2 kPa, the stiffness of healthy lung, up to 20 kPa, the stiffness of fibrotic foci in the lung(21).

**Equation 2** describes how collagen is secreted by fibroblasts and the daily degradation of collagen is integrated into the ABM:

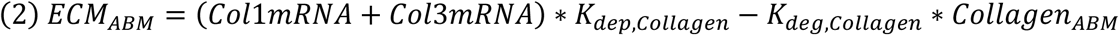

It is assumed that the ECM cannot be softer than that of observed human lung or stiffer than the observed fibrotic foci stiffness(21). Fibroblasts uptake TNF-α and IL-1β from the environment, and the relative activation of their receptors is calculated using the Hill Equation, although these cytokines are not explicitly modeled in the network model, as the pathway through which they regulate Thy-1 has not been described. Instead, we assume a 90% activation of both receptors is required for a Thy-1^+^ fibroblast to potentially transition and lose Thy-1 expression. **Equation 3** and **Equation 4** describe IL-1β uptake and TNF-α uptake, respectively:

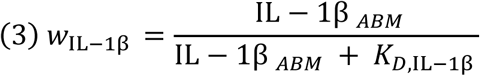

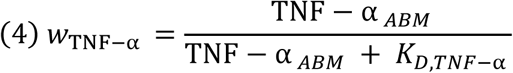

To account for uptake and degradation, up to 5% of the amount of cytokine on a patch was removed randomly every time a fibroblast agent encountered that patch. The network model also dictated fibroblast proliferation and apoptosis in conjunction with random chance. At every time step, the agent increased its age counter by one. The average lifespan was set to 55 days, with a deviation of 5 days which is similar to *in vitro* measurements of fibroblast lifespan(68). At each time step, there was a 25% chance a cell may undergo apoptosis as calculated by **Equation 5**, which integrates the network output for apoptosis for that cell with a stochastic factor based on the age of the cell:

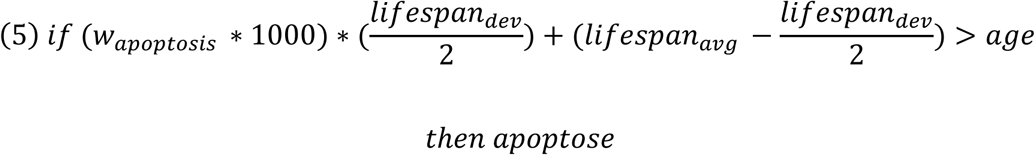

A cell proliferated according to the following equation that integrates network model output for proliferation with a stochastic in **Equation 6**:

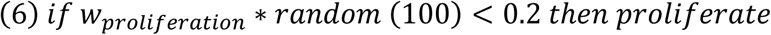

The ABM was simulated in NetLogo, and the logic-based network model was simulated in Python. These two models were connected via the “Py” extension in NetLogo, which allows each agent to call the network model and enter its specific inputs.

### Parameterization Using Machine Learning

Approximate Bayesian computation (ABC) is a method for generating the posterior probability distribution of a parameter. The parameter’s values are sampled from its prior distribution and fed into a model. Simulated data generated through ABC is then compared with observed data. The parameter values are retained only if the simulated data is sufficiently similar to the observed data. This process can be iterated so that parameter values that yield simulated data sets that better capture variability in observed data are retained more often than those that yield more disparate simulated data. In this way, a posterior probability distribution for the parameter is generated⁵¹. To generate simulated data with the NetLogo model, we used the Python package nl4py⁵² to interact with NetLogo.

A previously published dataset containing 36 patient lung biopsies was used as a validation dataset. These biopsies were scored based on the percent of fibrosis within a region of interest, with 1 being the lowest and 3 being the highest. To parameterize our model, we first identified a distribution of values for the probability of a fibroblast entering the alveolar space, which yielded a distribution of fibrosis scores similar to that of the validation data set. We assume that a fibroblast agent would only enter the alveolar space at points of alveolar barrier disruption (e.g., a fibrotic foci). A Thy-1⁻ fibroblast would enter the alveolar space if the sum of the ECM stiffnesses of the patch it is on, and four neighboring patches (north, east, west, and south of the agent) was greater than or equal to the variable “entry-threshold”. The maximum stiffness value for a patch was 1, therefore the maximum sum of the five patches sampled was 5. One hundred values of the entry-threshold were sampled from a flat beta prior distribution ranging from 0 to 5. For each sampled value, the NetLogo simulation was run 36 times to match the size of the validation data set. At the end of the simulations, the fibrosis score was recorded for three different possible end points: 365, 730, and 1095 time steps. To determine whether a sampled value should be retained, the algorithm used a chi-square test to compare the distribution of fibrosis scores in the validation set with that in the simulated data set. If no significant difference between the validation and simulated distributions was detected (P>0.05), that sampled value was retained. There were no significant differences in fibrotic foci formation based on the sampling of this variable alone, indicating that the model was not sensitive to the fibroblasts’ ability to enter the alveolar space. We then shifted our focus to another set of variables that the model might be sensitive to: the collagen deposition rate (K_dep_) and the collagen degradation rate (K_deg_).

For this second parameterization, three parameters were jointly optimized: K_dep_, K_deg_, and the Thy-1⁺ to Thy-1⁻ transition threshold. Rather than simple random sampling, we employed Bayesian optimization using the Python package scikit-optimize. ABC was used to iteratively optimize parameter combinations to minimize variability to control data. The search space was bounded as follows: K_dep_ ∈ [0.0, 200.0], K_deg_ ∈ [0.0, 200.0], and “transition-threshold” ∈ [0.0, 0.2]. All other simulation parameters were held fixed, including the initial counts of Thy-1⁺ and Thy-1⁻ fibroblasts (90 and 10, respectively), TNF and IL-1β thresholds (both set to 50), TGF-β (700 pg/mL), and the alveolar space entry threshold (4). For each candidate parameter set, 24 parallel simulations were run for 365 ticks (one simulated year), and fibrosis scores were collected at time steps of 365. The cost function minimized was the chi-squared statistic comparing the simulated distribution of fibrosis scores to the clinical validation distribution; this allowed the optimizer to quantify how closely each parameter set matched the observed data and to use that information to guide subsequent proposals. This process was repeated for 500 iterations using an Extra Trees surrogate model via the forest_minimize function. Partial dependence plots of the objective function landscape were generated to visualize the sensitivity of the fibrosis score distribution to each parameter and their pairwise interactions (**Fig 10**).

**Fig 10.**
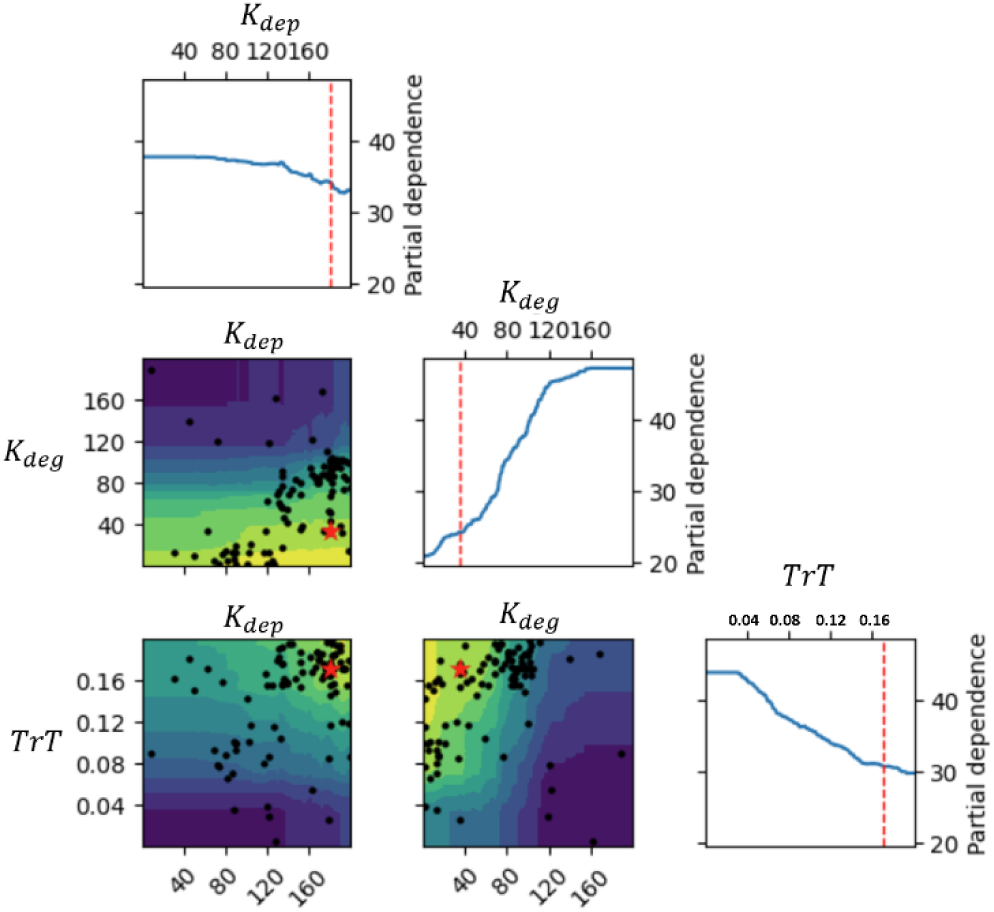
Parameterization of rate of collagen deposition and degradation. Correlation between K_deg_ and K_dep_, suggesting that if the relationship K_deg_ <= K_dep_ – 40, the model validates with experimental data.

### In vitro Fibroblast Culture

Commercially available human lung fibroblasts (ATCC, CCL-210) were expanded to passage 4. Collagen solution from bovine skin (Sigma-Aldrich, C4243) was prepared by diluting to 1 mg/mL in DPBS and pH-balancing with 1 N NaOH and HCl to a pH of between 7.2 and 7.4 (Day-2). As appropriate, 75 µL of collagen solution was added to each well of a 96-well tissue culture plate (Corning, 3603). The collagen solution was incubated at 37°C for 1 hour to allow solidification. Fibroblasts were trypsinized using 0.05% trypsin/EDTA (Gibco, 25300-054) and resuspended in standard cell culture media consisting of DMEM (Gibco, 11965-092), 10% fetal bovine serum (Gibco, 16000-044), and 1% penicillin-streptomycin (Gibco, 15140-122). Fibroblasts were seeded at 12,500 cells/cm^2^ (4,000 cells per well) in 200 µL of cell culture media on top of tissue culture plastic or formed collagen gel. The following day (Day-1), cell culture media was aspirated and replaced with 200 µL starvation media, consisting of DMEM (Gibco, 11965-092), 0.1% BSA (Sigma-Aldrich, A8412), and 1% penicillin-streptomycin. The following day (Day 0), starvation media was aspirated and replaced with 200 µL of fresh starvation media, supplemented with 50 ng/mL IL-1β (Biolegend, 579402) and 50 ng/mL TNF-α (Biolegend, 570102), as appropriate. Cells were incubated for 72 hours at 37°C, 5% CO_2_. Following incubation (Day 3), supernatant was collected, and cells were either fixed in 4% paraformaldehyde or lysed for later RNA analysis. Supernatant and lysate were stored at-80°C and fixed cells were stored at 4°C.

### Immunofluorescent Staining and Imaging of Cells

Cells were fixed in 100 µL of 4% paraformaldehyde (PFA) for 15 minutes (on shaker, in dark, at room temperature). PFA was removed, and wells were washed three times in 200 µL of DPBS, with a 5-minute incubation between washes (on shaker, at room temperature). Cells were permeabilized and blocked in 100 µL of perm/block buffer for 1 hour (on a shaker at room temperature). Perm/block buffer consisted of 0.3% Triton X-100 and 5% goat or donkey serum (as appropriate) in DPBS. Primary antibodies were prepared during this incubation. All antibodies (primary and secondary) were diluted in antibody dilution buffer containing 0.3% Triton X-100 and 0.1% BSA in DPBS. Primary antibodies used across the experiment are as follows: Mouse anti-human αSMA (Invitrogen, 14-9760-82, 1:200), mouse anti-human Col1a1 (Cell Signaling Technologies, 66948, 1:100), goat anti-human Col3a1 (Invitrogen, PA5-34787, 1:200), rabbit anti-human Thy-1 (Invitrogen, MA5-32559, 1: 100), and phalloidin conjugated with Alexa Fluor™ 546 (Invitrogen, A22283, 1:400). Following perm/block incubation, solution was aspirated and replaced with 100 µL of primary antibody dilution and incubated overnight (on shaker, in the dark, at 4°C).

The next day, the primary antibody dilution was removed, and the wells were washed three times in 200 µL of DPBS, with a 5-minute incubation between washes (on a shaker, in the dark, at room temperature). Secondary antibodies were prepared in antibody dilution buffer. Secondary antibodies used across the experiment are as follows: Goat anti-mouse Alexa Fluor 488 (Invitrogen, A-11001, 1:100) and donkey anti-rabbit Alexa Fluor 647 (Invitrogen, A-31573, 1:1000). Following washes, 100 µL of diluted secondary antibodies were added to appropriate wells, and incubated for two hours (on shaker, in the dark, at room temperature). After incubation, secondary antibodies were aspirated and wells were washed three times in 200 µL of DPBS, with a 5-minute incubation between washes (on shaker, in the dark, at room temperature). DAPI was prepared in DPBS (Invitrogen, R37606, 2 drops per mL), and 100 µL were incubated in the appropriate wells for 15 minutes (on a shaker, in the dark, at room temperature). The plates were immediately imaged on an Operetta CLS (Perkin Elmer).

Images were collected using the Operetta’s 10x air objective. Images were taken at the center of each well. For cells embedded in gel, Z-stack images were collected, and subsequent analysis was conducted on the max projection of these stacks. For image quantification, Integrated Optical Density (IOD) was calculated for each well. IOD is calculated by multiplying the fluorescent intensity by the fluorescent area, then dividing by the number of nuclei identified in each image region.

### qPCR

Following removal of supernatant, RNA was lysed in 100 µL of Buffer RLT (Qiagen, 79216). Lysate was stored at-80°C prior to isolation. Lysate was subsequently thawed and replicate wells were pooled. Pooled lysate was isolated using a commercially available kit (Qiagen RNeasy Mini Kit, 74104). Isolated RNA was quantified for concentration and protein contamination (Thermo Scientific NanoDrop™ OneC). RNA concentrations were normalized across samples with distilled water. Normalized RNA was combined with commercially available reagents (Invitrogen SuperScript™ VILO™ cDNA Synthesis Kit, 11754-050) and underwent reverse transcription polymerase chain reaction (RT-PCR) to create complementary DNA (cDNA). cDNA was combined with TaqMan™ Fast Advanced Master Mix (Applied Biosystems, 4444557) and TaqMan™ gene expression probes (**Table 2**). Samples underwent quantitative PCR (qPCR) reaction on a Bio-Rad CFX96 instrument. Quantification cycle (Cq) for each gene was normalized to HPRT1 (ΔCq). Relative expression for each gene was then calculated and reported.

**Table 2.** List of TaqMan expression probes.

|  |  |
| --- | --- |
| GUSB | Hs00939627_m1 |
| HPRT1 | Hs02800695_m1 |
| Col1a1 | Hs00164004_m1 |
| Col3a1 | Hs00943809_m1 |
| ACTA2 | Hs00426835_g1 |

### Histochemical Staining and Imaging of Patient Samples

Lung tissue samples that were previously collected, fixed in paraformaldehyde, and deidentified by the University of Chicago Medical Center were sectioned into 10 μm thick slices then mounted for histological analysis. A key matching diagnosis of IPF or non-IPF was also provided to the investigator performing the histological analysis. A variation of Movat’s Pentachrome staining was performed, consisting of only Verhoeff’s Elastic, Alcian Blue, and Alcoholic Saffron to resolve elastic fibers and nuclei (black), mucins and glycosaminoglycans (blue), and collagen (yellow), respectively. All samples in the same experiment were imaged using the 10X objective and K3C color camera on the Leica DMI8. To aid in visualization and analysis, the red and blue color channels were swapped, making collagen a bright blue. Images were then provided to another investigator who was blind to each sample’s diagnosis to train the Qupath algorithm on the collagen stain (blue). Once the pixel classifier was trained, 20 regions of interest (ROIs) were randomly generated, each the size of the computational viewer, on each tissue image. ROIs that contained an edge or a large vessel, which contains large amounts of collagen in the basement membrane, were removed, and a new random ROI was generated in their place. The average collagen content in each ROI was calculated as the number of blue pixels divided by the total number of pixels. ROIs for each patient sample were then averaged to generate the final collagen content for each patient. Once the analysis was complete, the sample diagnosis classifiers were revealed for data visualization.

### Computational Output Image Analysis using QuPath

At the six-month and twelve-month timepoints in the simulation, the model exported a greyscale image of the fibrotic foci formed to that point. These images were then loaded into QuPath, an open-source software designed for biological image data analysis(69). First, a training dataset was generated to calibrate the “Threshold” tool in QuPath to properly estimate fibrotic foci size and exclude foci that were only a single patch (10 microns or less). An automated image analysis pipeline was then constructed in QuPath’s environment to quantify the size and brightness characteristics for each foci in an image. Then a second dataset was generated that contained twenty images per time point per scenario and was uploaded into QuPath and the automated pipeline was run. Data was then uploaded to Excel, and foci characteristics were then averaged for all foci in an image. This data was then transferred to GraphPad Prism for principal component analysis and Pearson’s correlation analysis. The data used for this section of the chapter were from an “abbreviated” version of the multiscale model, due to the long computation time required to generate each data point from the full model. This abbreviated still contained the same rules and simplified the rate of collagen deposition that would normally be an output of the network model to specific values correlating with the ECM stiffness input that results in that collagen output.

### Statistics

For the experimental results, unpaired t-tests were used to compare the stimulated and unstimulated treatment groups. Two-way ANOVA followed by a multiple comparisons procedure (e.g., Tukey’s HSD) will be used to determine significant differences (asserting p<0.05) across samples of different starting Thy-1^-^ fibroblast populations and transition thresholds. When only one of these variables is different between groups, a one-way ANOVA followed by a multiple comparisons procedure (e.g., Tukey’s HSD) will be used to determine significant differences (asserting p<0.05) across samples.

## Acknowledgements

The authors would like to thank Dr. William Basener for his help in completing the approximate Bayesian computation portions of this work.

## Author Contributions

Figure and manuscript preparation: JLD, DJC, and SMP. Conceptualization: JLD, THB, CA, JJS, and SMP. Study design: JLD, THB, JJS, and SMP. Data collection: JLD, DJC, RTH, CS, DH, MB, RA, MA, TGE, TEV, AS, JMS, and DA. Data analysis: JLD, DJC, CS, MB, RA, MA, DH, and TGE. All authors participated in the review and approval of the final manuscript.

